# Netrin-1 directs vascular patterning and maturity in the developing kidney

**DOI:** 10.1101/2023.04.14.536975

**Authors:** Samuel Emery Honeycutt, Pierre-Emmanuel Yoann N’Guetta, Deanna Marie Hardesty, Yubin Xiong, Shamus Luke Cooper, Lori Lynn O’Brien

## Abstract

Blood filtering by the kidney requires the establishment of an intricate vascular system that works to support body fluid and organ homeostasis. Despite these critical roles, little is known about how vascular architecture is established during kidney development. More specifically, how signals from the kidney influence vessel maturity and patterning remains poorly understood. Netrin-1 (*Ntn1*) is a secreted ligand critical for vessel and neuronal guidance. Here, we demonstrate that *Ntn1* is expressed by stromal progenitors in the developing kidney, and conditional deletion of *Ntn1* from Foxd1+ stromal progenitors (*Foxd1^GC/+^;Ntn1^fl/fl^*) results in hypoplastic kidneys that display extended nephrogenesis. Despite expression of the netrin-1 receptor *Unc5c* in the adjacent nephron progenitor niche, *Unc5c* knockout kidneys develop normally. The netrin-1 receptor *Unc5b* is expressed by embryonic kidney endothelium and therefore we interrogated the vascular networks of *Foxd1^GC/+^;Ntn1^fl/fl^* kidneys. Wholemount, 3D analyses revealed the loss of a predictable vascular pattern in mutant kidneys. As vascular patterning has been linked to vessel maturity, we investigated arterialization in these mutants. Quantification of the CD31+ endothelium at E15.5 revealed no differences in metrics such as the number of branches or branch points, whereas the arterial vascular smooth muscle metrics were significantly reduced at both E15.5 and P0. In support of these results, whole kidney RNA-seq showed upregulation of angiogenic programs and downregulation of muscle-related programs which included smooth muscle-associated genes. Together, our findings highlight the significance of netrin-1 to proper vascularization and kidney development.

## Introduction

The organization of blood vessels into a network capable of maintaining organ and body fluid homeostasis requires both proper development of the vasculature and specialization unique to the tissue. Inappropriate vascular formation and patterning can result in pathologies in many organs(Blei and Bittman, 2016; Heimann and Damato, 2010; Queisser et al., 2021; Wälchli et al., 2023). Vascular endothelial cells organize into branched, hierarchical networks that form a continuous system for blood flow. Larger arteries branch into smaller arterioles, which further divide into capillaries. Venules then return the blood to larger veins and continue its circulation(Potente and Mäkinen, 2017). The vessels within these networks are distinct in their morphology and function, which is often tailored to the precise needs of the organ. For example, the endothelium of the kidney glomerular capillaries and ascending vasa recta are fenestrated to enable their unique functions in filtration and diffusion, respectively(Molema and Aird, 2012; Pallone et al., 2003). In association with their functions, the endothelium of these different vascular beds also have distinct molecular signatures(Barry et al., 2019; Daniel et al., 2018). Additionally, endothelial association with mural cells delineates the various types of vessels within an organ. Larger vessels such as arteries are coated by vascular smooth muscle cells whereas smaller capillaries are covered by pericytes, with each type of mural cell supporting distinct vascular functions(Potente and Mäkinen, 2017). Altogether, these vascular properties highlight the need for precise developmental programs that support the proper formation of these complex networks.

Recent studies have begun to shed light on the progressive formation of kidney vascular networks during development. The temporal progression and predictable rudimentary patterning of the major renal arteries through mid-gestation were elucidated through comprehensive imaging studies(Daniel et al., 2018; Munro et al., 2017). Arteries which connect to the aorta undergo angiogenic growth towards the developing kidney forming a vascular ring around the base of the ureteric bud by E11.25(Munro et al., 2017). Continued growth, remodeling, and maturation of the endothelial network leads to the establishment of an arterial tree by E13.5 that forms a predictable pattern(Daniel et al., 2018). Renin expressing cells are localized along the developing arterial vasculature and mark points of branching, and renin itself is important for proper vascular development (Gomez et al., 1989; Reddi et al., 1998; Sequeira-Lopez et al., 2015b; Takahashi et al., 2005). At the molecular level, vascular patterning was shown to be regulated by the association of mural cells with the nascent endothelium and modulated in part by Pbx1(Hurtado et al., 2015). Deletion of the transcription factor *Pbx1* from the Foxd1+ stromal progenitors induces precocious differentiation and premature association of vascular mural cells with the endothelium, leading to stochastic arterial patterning and impaired renal function(Hurtado et al., 2015). Another study showed that Dicer activity in the kidney stroma is important for patterning smaller vessels, such as the peritubular and glomerular capillaries(Nakagawa et al., 2015). While these studies have garnered some of the first insights into how the kidney vascular network is established and organized, we still lack a clear understanding of how this process is modulated at the molecular level.

Signaling molecules play important roles during development, often providing instructional cues which help pattern the developing tissues. In respect to vasculature, a number of signaling pathways, which also regulate axon guidance, provide angiogenic guidance. This includes Slit/Robo, Eph/Ephrin, Semaphorins/Plexin-Neuropilins, and Netrins/UNC5(Adams and Eichmann, 2010). These guidance cues direct vascular development during embryogenesis, and unsurprisingly, the removal of many of these cues or their receptors results in vascular defects and mortality in the forming embryo (Bin et al., 2015; Gerety et al., 1999; Gitler et al., 2004; Gu et al., 2005, 2003; Jones et al., 2008; Lu et al., 2004). Netrin-1 (*Ntn1*) is a classical neuronal guidance cue well-known for its role in commissural axon guidance(Kennedy et al., 1994; Serafini et al., 1994). However, roles for netrin-1 have expanded to include mediating pancreas development, inner ear morphogenesis, mammary gland development, lung branching, and angiogenesis (Larrivée et al., 2007; Liu et al., 2004; Lu et al., 2004; Salminen et al., 2000; Srinivasan et al., 2003; Sun et al., 2011; Wilson et al., 2006; Yebra et al., 2003). Endothelial guidance by netrin-1 is accomplished through chemotactic interactions with its cognate receptors in the vasculature, Unc5b and CD146 (*Mcam1*), which regulate blood vessel growth, branching, and patterning by promoting either a repulsive or attractive response, respectively (Larrivée et al., 2007; Lu et al., 2004; Tu et al., 2015). Whether netrin-1 plays a similar role in guiding vascular development and patterning in the kidney remains unexplored.

Here we report that netrin-1 (*Ntn1*) is expressed and secreted by stromal progenitors in the developing kidney. Deletion of *Ntn1* from this population results in hypoplastic kidneys and delayed cessation of nephrogenesis. In correlation with its known role in vascular development, we found that netrin-1 supports proper patterning and maturation of the vasculature during kidney organogenesis. Our findings correlate with those in the accompanying manuscript by **Luo, Gu et al.**, and together we establish a critical role for netrin-1 in mediating proper kidney development through regulation of vascular maturity.

## Results

### Ntn1 is expressed and secreted by Foxd1+ stromal progenitors in the developing kidney

While netrin family members including netrin-1 play important roles in other branching organs during development, such as the lung, pancreas, and mammary gland, whether they have a role in kidney development has not been investigated to date(Liu et al., 2004; Srinivasan et al., 2003; Yebra et al., 2003). To begin, we examined the expression pattern of *Ntn1* mRNA in the developing kidney and found that *Ntn1* is expressed by stromal progenitors around the nephron progenitor caps at embryonic day (E)15.5 and, to a lesser extent, in the proximal tubule which is more specifically detected at later postnatal (P2) stages of development (Fig. 1A; Fig. S1A). This cellular expression pattern is confirmed by publicly available single cell RNA-seq data of the kidney at E18.5(Combes et al., 2019; England et al., 2020). As early as E11.5, secreted netrin-1 protein was observed surrounding the Six2+ nephron progenitors where the stromal progenitors coalesce (Fig. 1B). At E15.5, netrin-1 was similarly secreted into the area surrounding the nephron progenitor niche (Fig. 1C, arrow). Some protein was diffused into the area below the nephron progenitor niche where ureteric tips lie, potentially acting as a ligand sink to prevent further diffusion (Fig. 1C). The stromal progenitors expressing *Ntn1* give rise to the renal interstitium, mesangial cells, and vascular mural cells, among others, and function in advancing nephron progenitor differentiation(Hatini et al., 1996; Kobayashi et al., 2014; Sequeira-Lopez et al., 2015a), suggesting that netrin-1 signaling may support proper kidney development.

**Figure 1.**
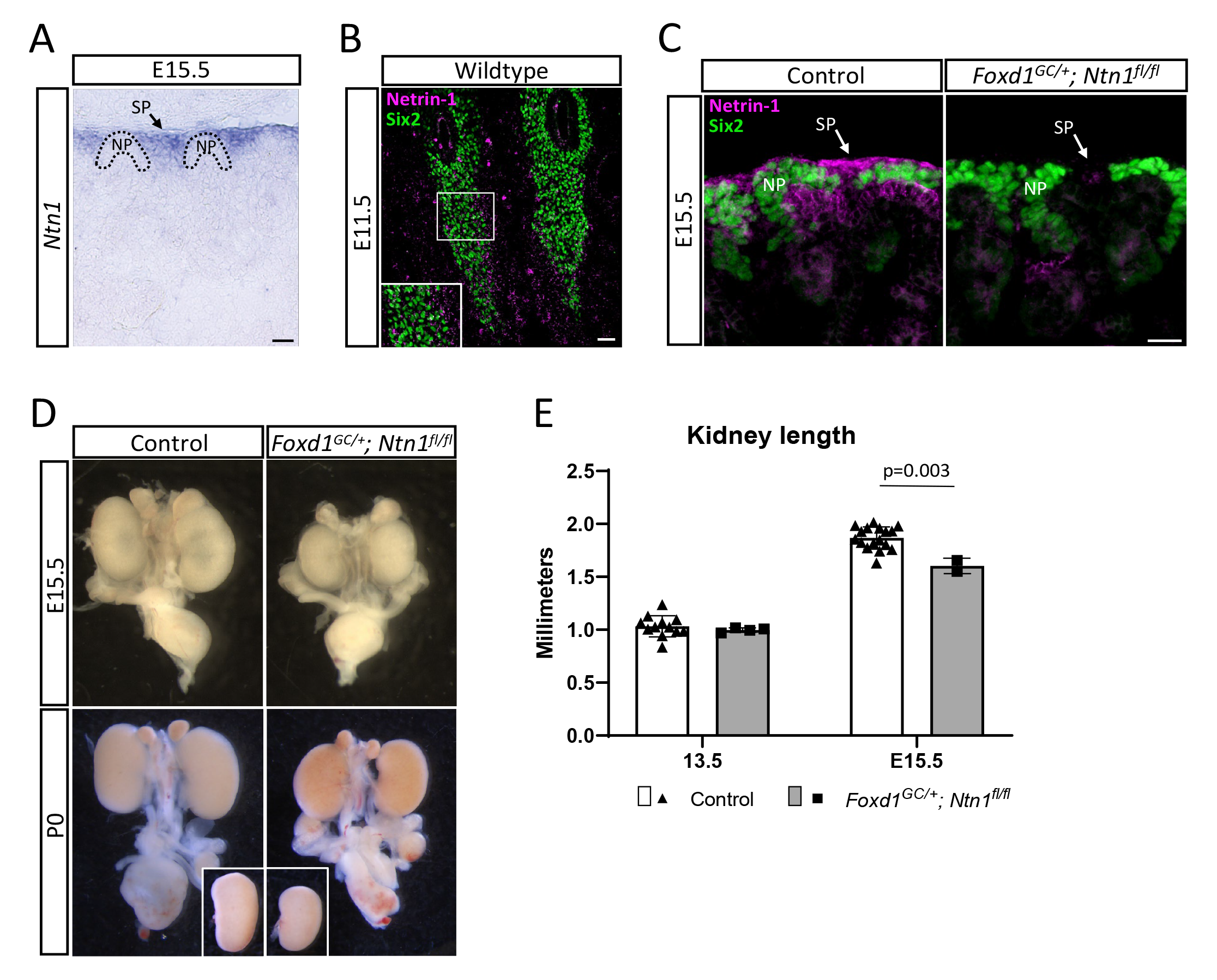
*Ntn1* is expressed and secreted by stromal progenitors in the kidney and deletion results in kidney hypoplasia. (A) Light microscopy image of an E15.5 section showing the *in situ* hybridization expression pattern of *Ntn1*. *Ntn1* expression is observed in the stromal progenitors (SP) surrounding the nephron progenitors (NP). Dotted lines demarcate NP niches. Scale bar=50 μm. (B) Immunostaining of E11.5 kidney sections shows that netrin-1 (magenta) is secreted by the stromal progenitors surrounding the Six2+ (green) nephron progenitors. Inset shows a higher magnification of the netrin-1 signal adjacent to the nephron progenitors. Scale bar=50 μm. (C) Immunofluorescence images showing netrin-1 (magenta) signal in the stromal progenitors (SP) and adjacent to the Six2+ nephron progenitors (NP, green) (left panel). Secreted netrin-1 signal is also observed in the ureteric tip region. Genetic deletion of *Ntn1* in the *Foxd1*+ stromal population results in a significant reduction in netrin-1 signal (right panel). Scale bar=25 μm. (D) Kidney hypoplasia is observed at various stages of development. Shown are micrographs of whole urogenital systems at E15.5 and P0. Insets for P0 show the same kidneys micro dissected and side-by-side for size comparison. (E) Kidney length was measured in Fiji imaging software at E13.5 and E15.5, with the size reduction becoming statistically significant by E15.5. Columns and error represent mean and standard deviation (SD), respectively, with significance derived from Students’ T-test. Respective p-values displayed on graphs.

### Deletion of Ntn1 from stromal progenitors disrupts normal kidney development

Germline deletion of *Ntn1* was previously shown to result in embryonic lethality by E14.5(Bin et al., 2015). To circumvent this early lethality and address the role of netrin-1 in kidney development, we conditionally deleted *Ntn1* from the Foxd1+ stromal progenitors by using the *Foxd1^GC^* allele in combination with a *Ntn1* floxed allele(Bin et al., 2015; Humphreys et al., 2010; Kobayashi et al., 2014). The resulting *Foxd1^GC/+^; Ntn1 ^fl/fl^* mice have very little or no netrin-1 protein present in the interstitial regions surrounding the nephron progenitors, confirming efficient deletion (Fig. 1C). Our initial observations found that *Foxd1^GC/+^; Ntn1 ^fl/fl^* mutant animals have hypoplastic kidneys at E15.5 and this phenotype persists postnatally (Fig. 1D). Immunostaining of the ureteric tree at E14.5 also revealed qualitative differences in kidney size (Fig. S1B). Measurements of kidney length showed these reductions became most significant after E13.5 (Fig. 1E).

As a means of assessing whether *Ntn1* deletion affects either the onset of kidney development or cessation of nephrogenesis, we examined the presence of Six2+ nephron progenitors at the initial branching event (E11.5) and after the cessation of nephrogenesis (P4), which normally occurs around P2-P3 in the mouse(Hartman et al., 2007; Self et al., 2006). At E11.5, we observed no empirical differences in the condensation of Six2+ nephron progenitors around the ureteric buds suggesting kidney development initiated at the same time in *Ntn1* conditional mutants (Fig. 2A). Analysis of nephron progenitor niches at E15.5 revealed no obvious differences in their organization, suggesting normal progression of nephrogenesis (Fig. 2B). Strikingly though, at P4, *Foxd1^GC/+^; Ntn1^fl/fl^* kidneys maintained small clusters of Six2+ cells whereas controls had exhausted their Six2+ population as expected (Fig. 2C). Our previous bulk nephron progenitor RNA-seq data as well as recent scRNA-seq data indicate the netrin receptor *Unc5c* is expressed by nephron progenitors(Combes et al., 2019; O’Brien et al., 2018, 2016). *In situ* hybridization of *Unc5c* confirmed expression by the nephron progenitors, as well as podocytes and smooth muscle cells of the ureter (Fig. 2D). Due to the nephron progenitor expression, we wondered whether disrupted signaling via netrin-1 to the *Unc5c+* nephron progenitors might account for the smaller kidney size and delay in cessation of nephrogenesis. We assessed the kidney phenotype of whole animal *Unc5c* knockouts(Ackerman et al., 1997) and found no difference in kidney size compared to controls (Fig. 2E). Additionally, the Six2+ nephron progenitor niches appear normal, and no other overt phenotypes were observed at any stages examined (Fig 2F). Therefore, the absence of netrin-1 signaling through Unc5c is unlikely to cause the observed hypoplastic phenotype.

**Figure 2.**
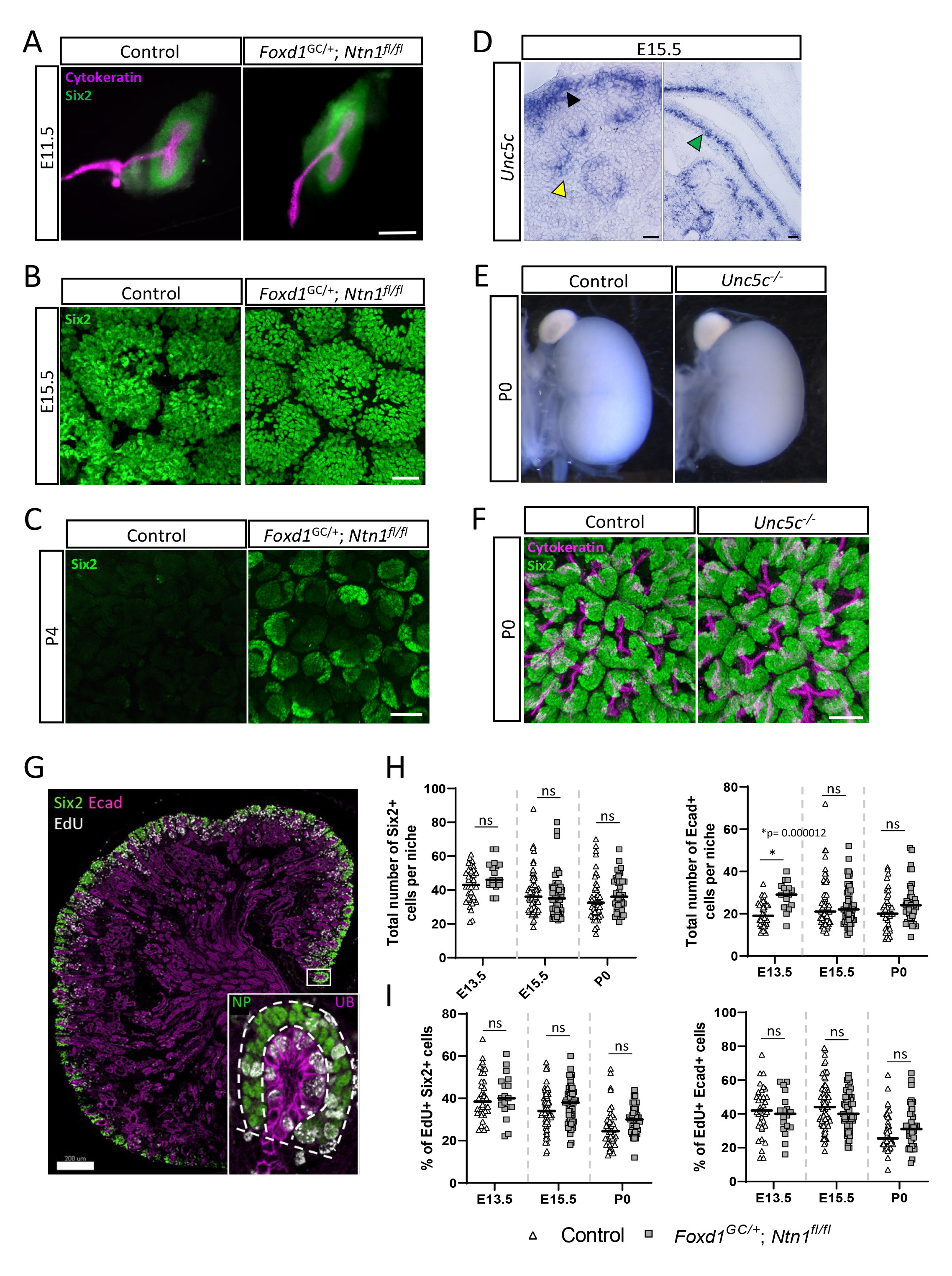
Loss of *Ntn1* delays the cessation of nephrogenesis but is not due to signaling through nephron progenitor expressed *Unc5c* or differences in progenitor dynamics. A) Immunostaining of whole metanephric kidneys at E11.5 shows that metanephric kidney development is initiated normally in *Ntn1* conditional knockouts, with the expected T-shaped ureteric bud (cytokeratin, magenta) surrounded by condensed nephron progenitors (Six2, green). Scale bar=200 μm. B) Nephron progenitor niches (Six2, green) appear normally organized at E15.5 in surface views of wholemount immunostained kidneys. Scale bar=30 μm. C) Wholemount immunostains of P4 kidneys show the persistence of Six2+ nephron progenitors (green) on the surface of the kidney in *Ntn1* conditional knockouts. Scale bar=30 μm. D) *In situ* hybridization showing the expression of the netrin-1 receptor *Unc5c* in the nephron progenitors (black arrowhead), podocytes (yellow arrowhead), and smooth muscle layer of the ureter (green arrowhead) at E15.5. Scale bar=50 μm. E) Light microscopy images of whole kidneys at P0 showing no difference in kidney size of *Unc5c* knockout kidneys compared to controls. F) Wholemount immunostaining and surface views show the normal organization of nephron progenitor niches in *Unc5c* knockouts at P0 (Six2-green, cytokeratin-magenta). Scale bar=40 μm. G) Example of a sectioned kidney labeled with EdU (white) and immunostained with Six2 (green) and E-cadherin (Ecad, magenta) to label the nephron progenitors and collecting duct system, respectively. Inset shows a high magnification image of a niche with dotted lines highlighting the extent of what was considered either a nephron progenitor (NP) niche or ureteric bud (UB) tip niche for quantification purposes. Scale bar=200 μm. H) Quantification of Six2+ nephron progenitors or Ecad+ ureteric tip cells at E13.5, E15.5, and P0 show no significant differences in total cell numbers except for Ecad+ cells at E13.5. Bars represent mean; significance was determined using Students t-test. I) Quantification of the % of EdU+ Six2+ nephron progenitors or EdU+ Ecad+ ureteric tip cells reveals no significant difference at E13.5, E15.5, or P0 between *Foxd1^GC/+^*; *Ntn1^fl/fl^* kidneys or controls. Bars represent mean; significance was determined using Students t-test.

In developing *Foxd1^GC/+^; Ntn1 ^fl/fl^* kidneys, the loss of *Ntn1* could affect the number of progenitors in nephron progenitor or ureteric tip niches, leading to hypoplasia. To assess cell numbers, we analyzed kidney sections at multiple timepoints and counted Six2+ nephron progenitors and Ecad+ ureteric tip cells within the niche (Fig. 2G). Apart from a transient difference in tip cell numbers at E13.5, neither population showed any significant differences between *Foxd1^GC/+^; Ntn1 ^fl/fl^* kidneys and controls when assessed at E13.5, E15.5, and P0 (Fig. 2H). Using the same method, we investigated whether the proliferative capacity of niche progenitors was impacted by measuring EdU incorporation into these populations as a measure of cell cycling (Fig. 2G). We found no significant differences in the number of EdU+ cells in either Six2+ or Ecad+ progenitor populations suggesting cell proliferation, at least at the stages analyzed, was not causing the hypoplastic phenotype (Fig. 3C).

**Figure 3.**
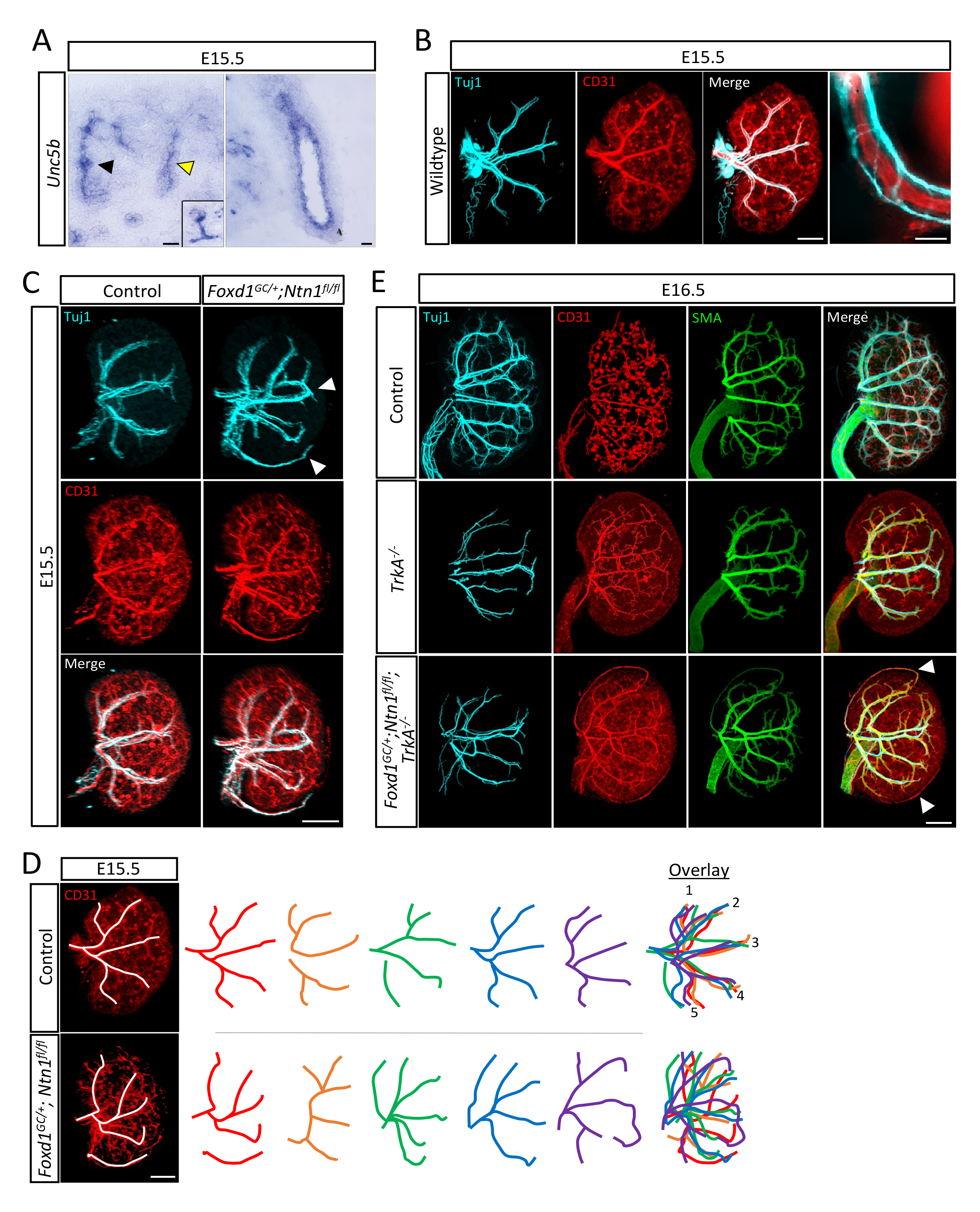
Stromal deletion of *Ntn1* results in aberrant neurovascular patterning. A) *In situ* hybridization on E15.5 kidney sections shows the expression of the netrin-1 receptor *Unc5b* in the endothelium of the kidney (black arrowhead and inset of glomerular endothelium, left panel), the secondary branches/generations of the collecting duct (yellow arrowhead, left panel), and ureter (right panel). Scale bars=50 μm B) Wholemount immunostain of an E15.5 wildtype kidney show the tracking of Tuj1+ (cyan) axonal projections with the renal vasculature (CD31, red). Far right panel shows an independent close-up image of a neurovascular bundle. Merge scale bar=300 μm, close-up scale bar=50 μm. C) Wholemount immunostains show the organization of the nerves and vasculature at E15.5 in *Nnt1* conditional knockouts and controls. The nerves (Tuj1, cyan) track with the endothelium (CD31, red) in all genotypes. White arrowheads highlight the abnormal patterning observed in *Ntn1* conditional mutants such as looping and ectopic neurovascular bundles. Scale bar=300 μm. D) Depiction of tracing (white lines) of the neurovascular bundles (left panel). Tracing of five control kidneys (top panel, colored lines) show predictable patterns of 5 main lateral arterial branches which can be overlayed. *Ntn1* conditional knockout kidneys (bottom panel, colored lines) show stochastic patterning with the absence of any predictable patterning. Scale bar=300 μm. E) Wholemount immunostains of control, *TrkA^-/-^*, and *Foxd1^GC/+^*; *Ntn1^fl/fl^; TrkA ^-/-^* kidneys at E16.5 show the organization of nerves (Tuj1, cyan), endothelium (CD31, red), and smooth muscle cells (SMA, green). Innervation of the kidney is reduced in *TrkA* knockouts, but abnormal patterning persists with vasculature not associated with axons (arrowheads) still showing abnormal patterning, suggesting the endothelium is the primary responder to netrin-1 guidance cues. Scale bar=300 μm.

### Netrin-1 mediates vascular patterning in the developing kidney

Both axons and endothelium respond to guidance cues such as netrin-1 using specific receptors. Unc5b has been shown to mediate angiogenesis (Larrivée et al., 2007; Lu et al., 2004) and *in situ* hybridization at E15.5 shows that *Unc5b* is expressed by the kidney endothelium as well as the ureteric epithelium, although is excluded from the most peripheral ureteric branches(Lindström et al., 2018) (Fig 3A). During kidney development, axonal projections tightly associate and track with the vasculature, forming tight neurovascular bundles (Fig 3B). Therefore, we included both nerves and vessels in our analyses since both have the potential to respond to netrin-1 guidance. To assess overall organization and patterning of these networks, we utilized wholemount immunostaining in combination with light-sheet microscopy to generate 3D visualizations. Control kidneys show the formation of an extensive CD31+ endothelial network by E15.5 (Fig. 3C). Tuj1+ axons track closely with the main arterial branches at this stage which form a pattern consistent with those observed by Daniel et al.(Daniel et al., 2018) (Fig 3C, Suppl. Video 1). In contrast, we observed aberrant neuronal and vascular patterning in the *Foxd1^GC/+^; Ntn1 ^fl/fl^* mutant kidneys (Fig. 3C, Suppl. Video 2). The example shown features looping of the arterial tree within the kidney cortex and ectopic vessels on the kidney surface (arrowheads, Fig. 3C). Tracing of the main CD31+ arterial branches shows that controls form a predictable pattern, as expected, which can be overlayed into 5 main branches from a lateral view(Daniel et al., 2018) (Fig. 3D). However, the *Foxd1^GC/+^; Ntn1^fl/fl^* kidneys have a breakdown of predictable patterning and overall stochastic patterning (Fig. 3D). Though not a focus of this study, the cortical vasculature around the developing nephron progenitor niches of *Foxd1^GC/+^; Ntn1 ^fl/fl^* kidneys is also disrupted, featuring disorganized and ectopic vasculature (Fig. S2A). Notably, the presence of Ter119+ erythroid cells indicates that the blood vessels are perfused in both control and *Foxd1^GC/+^; Ntn1^fl/fl^* kidneys at E15.5, although contribution from hemo-vasculogenesis is also possible (Sequeira Lopez et al., 2003)(Fig. S2B). *Ntn3* (netrin-3), a functionally related guidance cue that can engage netrin-1 receptors(Wang et al., 1999), is also expressed, albeit at lower levels, by the stromal progenitors(Combes et al., 2019; England et al., 2020) (Fig S3A). However, we did not observe any neurovascular patterning defects or differences in kidney size in *Ntn3* knockouts or any exacerbation of the *Foxd1^GC/+^;Ntn1^fl/fl^* phenotype when double knockouts (*Foxd1^GC/+^;Ntn1^fl/fl^;Ntn3^-/-^)* were generated (Fig. S3B,C). Together, these data suggest netrin-1 is the primary netrin guidance cue responsible for the neurovascular patterning defects.

As both axons and endothelium can respond to netrin-1 guidance cues, we sought to differentiate the primary responder to stromal-derived netrin-1 signals. Deletion of the *TrkA* receptor for Nerve Growth Factor (NGF) induces apoptosis of peripheral neurons such as those that innervate the kidney(Smeyne et al., 1994). We investigated vascular patterns in the setting of reduced kidney innervation utilizing a *TrkA* knockout allele(Liebl et al., 2000). In *TrkA* knockout kidneys, the vascular pattern was qualitatively normal at E16.5 (Fig. 3E). However, when combined with the *Foxd1^GC/+^;Ntn1^fl/fl^* alleles to generate *Foxd1^GC/+^;Ntn1^fl/fl^;TrkA^-/-^* animals, the kidney endothelium still displayed abnormal patterning including looping and ectopic vessels (Fig. 3E). The ectopic and looping vessels were not associated with Tuj1+ axons (Fig. 3E, arrowheads), suggesting the endothelium is the primary responder to netrin-1 guidance cues with the nerves following the vascular pattern, and in particular the smooth muscle coated (alpha-smooth muscle actin, SMA+) endothelium (Fig. 3E). We also wanted to investigate whether *Unc5c* knockout kidneys displayed altered patterning, even though they were not hypoplastic. The neurovascualture appeared largely normal with no ectopic vessels, aberrant looping, or qualitative reductions suggesting Unc5c is not a major receptor for netrin-1 in the kidney or is functionally redundant with another receptor (Fig. S3D).

### Vascular smooth muscle cell coverage is reduced in Foxd1^GC/+^; Ntn1^fl/fl^ kidneys

As the endothelium matures it is covered by a layer of vascular smooth muscle cells (vSMCs). These cells provide support, maintain arterial wall integrity, and can direct innervation through the release of guidance molecules(Brunet et al., 2014; Potente and Mäkinen, 2017). To determine if vSMCs were impacted in *Foxd1^GC/+^; Ntn1^fl/fl^* animals, we investigated SMA coverage of the endothelium in wholemount immunostained kidneys. Despite the lack of predictable patterning, quantification of the CD31+ endothelium in *Foxd1^GC/+^; Ntn1^fl/fl^* kidneys revealed that branching metrics and total vessel length were not significantly affected at E15.5 (Fig S4A, Fig 4A). However, the CD31+ endothelium showed qualitative reductions in SMA coverage in *Foxd1^GC/+^; Ntn1 ^fl/fl^* kidneys compared to controls (Fig. 4B, arrowheads, Suppl. Videos 3,4). Furthermore, *Tuj1*+ axons tracked exclusively with SMA+ vSMCs, representing arteries, and were not found anywhere where endothelium singularly expressed CD31. These data suggest that *Ntn1* loss impacts smooth muscle coverage of the vasculature, which is associated with vessel stabilization and maturity.

**Figure 4.**
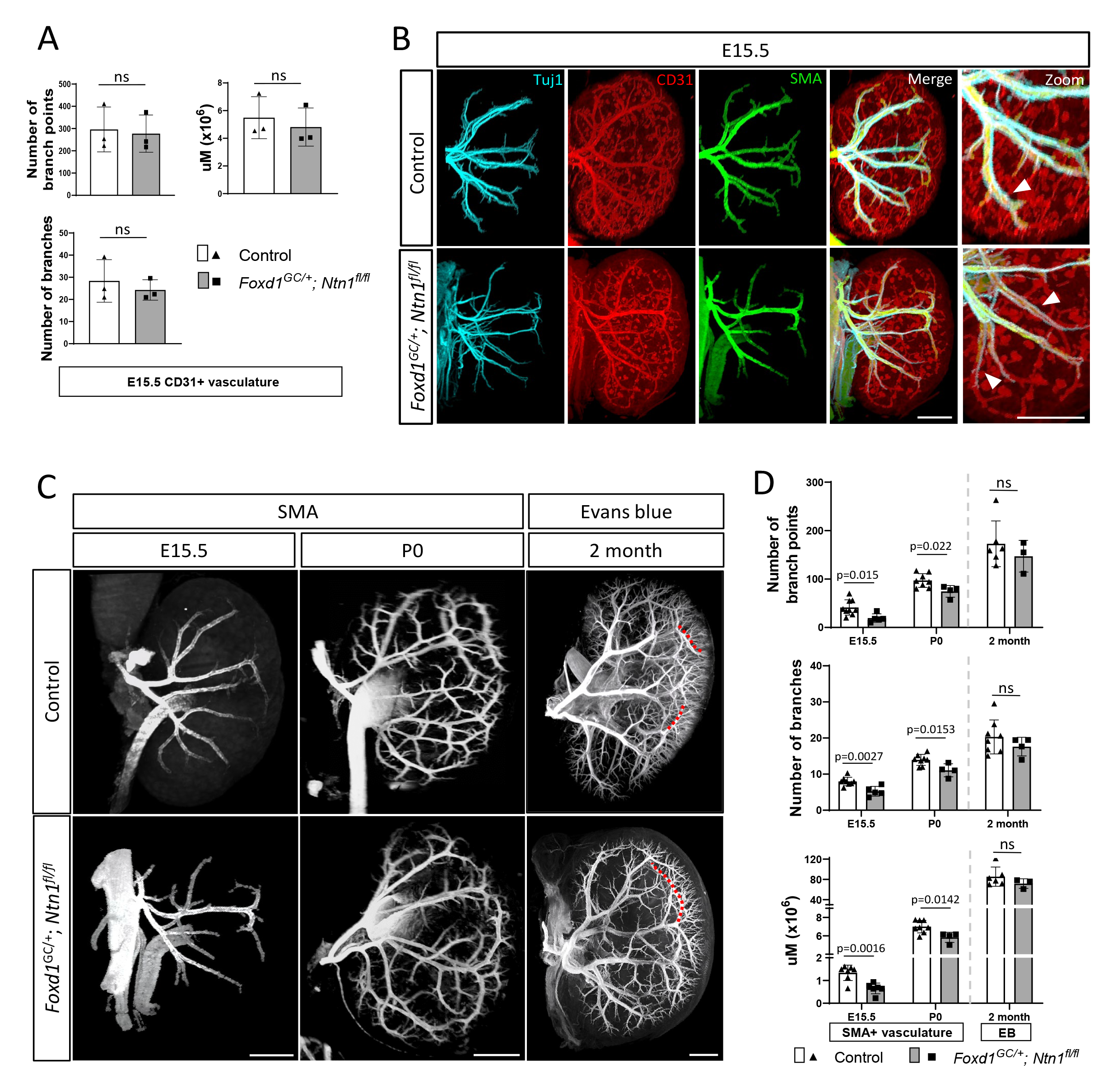
*Ntn1* conditional knockouts have normal endothelial metrics but reduced smooth muscle coverage throughout development. A) Quantitation of CD31+ endothelial metrics including the number of branches, branch points, and total endothelium (total additive length, μM) at E15.5 shows no statistical differences between controls and *Ntn1* conditional knockouts. Significance was calculated with Students t- test, p<0.05 was considered significant, columns represent mean. Error bars = SD. B) Wholemount immunostaining of E15.5 kidneys with smooth muscle actin (SMA, green) in addition to the axonal (Tuj1, cyan) and endothelial (CD31, red) markers show a qualitative delay in smooth muscle cell association with the endothelium (white arrowheads, zoom) suggesting a potential delay is vascular maturation. Scale bars=300 μm. C) Wholemount immunostains of E15.5 and P0 kidneys with SMA (white) show a qualitative reduction on SMA vascular coverage in *Ntn1* conditional mutants. The injection of Evans Blue dye at 2 months to label the adult kidney vasculature shows a well-developed arterial tree with some remnants of abnormal patterning such as elongated branches in *Ntn1* conditional knockouts. Red dotted lines overlay normal (top) and abnormal (bottom) branching. E15.5 scale bar=300 μm, P0 scale bar=500 μm, 2-month scale bar=1000 μm. D) Quantitation of SMA+ arterial metrics including the number of branches, branch points, and total vasculature length (μM) at E15.5 and P0 show a deficit in smooth muscle metrics suggesting a delay in arterial differentiation. Quantitation of Evans blue labeled arterial vascular metrics at 2 months reveals that *Ntn1* conditional knockouts have attained levels similar to controls. Significance was calculated with Students t-test, p<0.05 was considered significant, individual p-values are displayed on the respective graphs, while columns represent mean. Error bars = SD. Demarcation between P0 and two months denotes the differences in labeling, with early stages labeled by SMA and 2 months perfused with Evans blue.

While vascular patterning defects in *Foxd1^GC/+^; Ntn1 ^fl/fl^* kidneys are apparent, they are stochastic and highly variable. Since some kidneys appear to have a more significant degree of mis-patterning, they may also display differences in vSMC coverage. However, despite the stochastic patterning differences, quantification of the SMA+ arterial vasculature showed reductions at E15.5 and P0 in *Foxd1^GC/+^; Ntn1^fl/fl^* kidneys (Fig. 4C,D). On average, E15.5 kidneys from *Foxd1^GC/+^; Ntn1 ^fl/fl^* animals have greater than a 20% reduction in SMA+ arterial metrics including branch number, branch points, and the total length of the vSMC-lined vasculature (Fig. 4D). At P0, reductions in these three vascular parameters were still significant highlighting the persistence of this phenotype throughout development. To determine whether the defects observed in development extended into adulthood, we labelled the vasculature of 2-3 month old *Foxd1^GC/+^; Ntn1 ^fl/fl^* animals with Evans blue (Fig 4C). Evans blue is a bis-azo dye that fluoresces only while bound to albumin and labels the perfused renal vasculature(Honeycutt and O’Brien, 2021). At two months, quantification of *Foxd1^GC/+^; Ntn1 ^fl/fl^* kidneys labeled with Evans blue revealed no significant differences in vascular metrics, although all three metrics trended towards a reduction (Fig. 5D). At both 2 and 3 months of age, ectopic vessels were no longer observed and the kidney vasculature has fewer visible defects, although this is difficult to ascertain due to the highly complex vascular network that has formed. However, differences were noted such as meandering, elongated vessels near the cortex (Fig 4C and Fig S4B, red dotted lines) and areas of reduced branching (Fig S4B, yellow arrows). Additionally, we occasionally observed clouds of Evans blue signal outside the vascular network (Fig S4B, white arrowhead) indicating possible leakage and compromise to vessel integrity.

**Figure 5.**
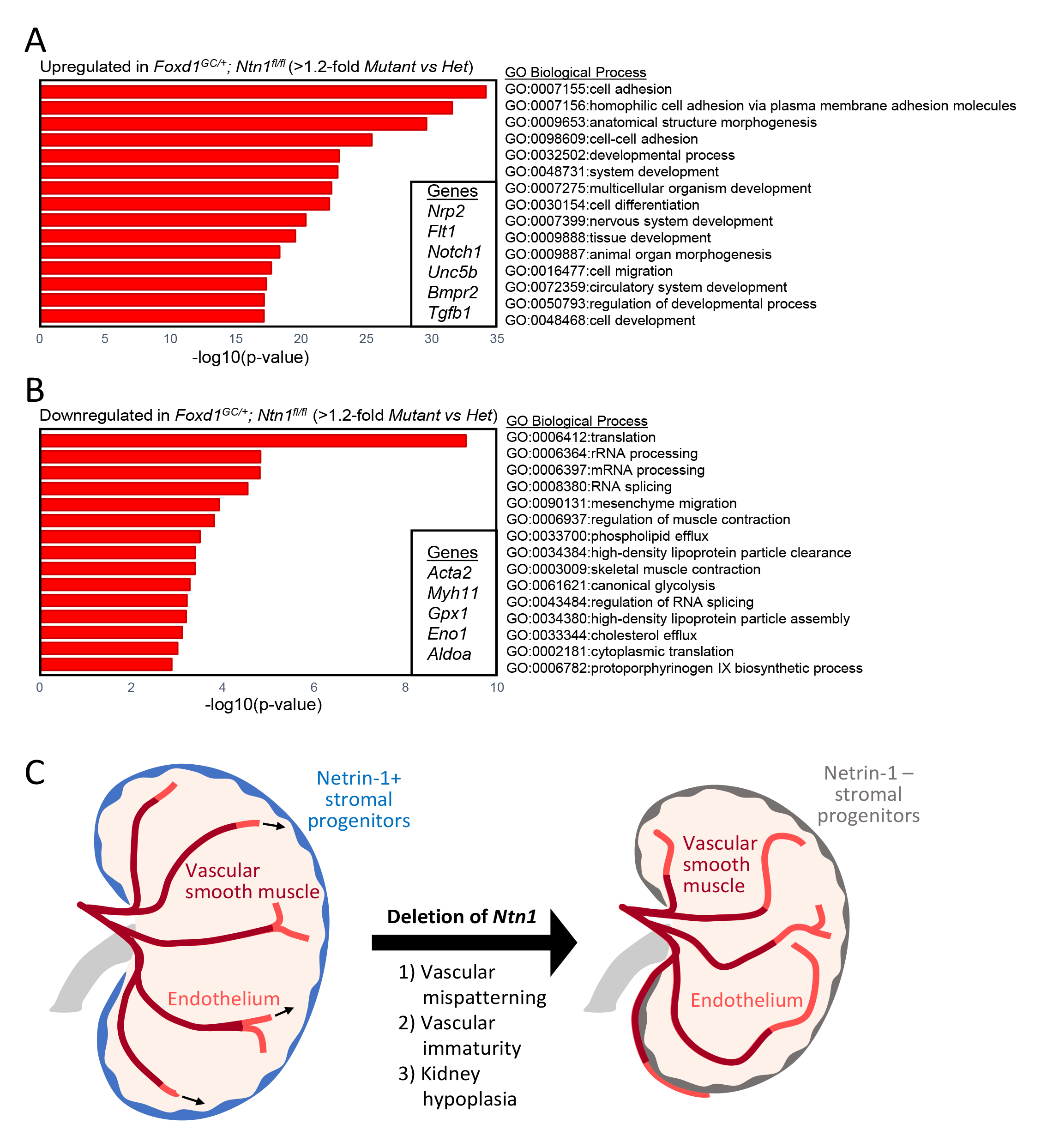
Gene expression analyses support a delay in vascular maturation in *Ntn1* conditional knockouts. A) Gene ontology (GO) analysis of genes upregulated (top) or B) downregulated (bottom) in *Ntn1* conditional knockouts. Shown are the GO Biological Processes enriched in both categories. Genes were considered up- or downregulated with a >1.2 fold-change. In the insets are examples of genes that are upregulated (top) or downregulated (bottom). C) Schematic of the role of netrin-1 in guiding vascular patterning and maturation during development.

### RNA-seq reveals differences in vascular and metabolic programs of Foxd1^GC/+^; Ntn1^fl/fl^ kidneys

To better understand alterations to transcriptional programs relevant to the vascular immaturity and the hypoplastic phenotype observed in *Foxd1^GC/+^; Ntn1^fl/fl^* animals, we performed bulk RNA-seq on whole kidneys (Table S1). To account for any exacerbated differences due to the loss of one *Foxd1* allele from the *GFP-Cre* insertion, we compared *Foxd1^GC/+^; Ntn1^fl/fl^* mutant kidneys to *Foxd1^GC/+^; Ntn1^fl/+^* heterozygous kidneys. We utilized a cutoff of 1.2-fold as more significant changes may be drowned out due to the bulk analysis. Gene ontology analyses were then performed on up- and downregulated genes to identify biological processes enriched in either category (Tables S2 and S3). In support of our imaging analyses, processes related to the circulatory system and blood vessel development were upregulated in *Foxd1^GC/+^; Ntn1^fl/fl^* kidneys and included genes such as *Notch1*, *Flt1*, *Nrp2*, and *Unc5b* (Fig. 5A, Table S2). Programs related to cell adhesion and cell migration were also upregulated and included genes in the cadherin superfamily and planar cell polarity related genes, respectively (Fig 5A, Table S2). Such pathways may affect the vSMC association with the endothelium in addition to affecting overall kidney development, thus leading to the hypoplastic phenotype. In support of the reduction in vSMC coverage of the endothelium in mutant kidneys, muscle related processes were downregulated and genes in these categories included *Acta2* which encodes for *α*-SMA and *Myh11* which is a smooth muscle myosin (Fig 5B, Table S3). Additional biological processes downregulated in *Foxd1^GC/+^; Ntn1^fl/fl^* kidneys were related to more general cellular processes such as translation, RNA processing, and glycolysis (Fig 5B). Altogether, these data suggest that in the absence of netrin-1 signaling, the kidney vasculature is more immature (increased angiogenic processes vs arterialization) which may affect metabolic and cellular processes, leading to the hypoplastic phenotype (Fig. 5C).

## Discussion

In this study, we demonstrate that stromal progenitor *Ntn1* is necessary for proper patterning and maturation of the kidney arterial network. *Foxd1^GC/+^; Ntn1 ^fl/fl^* kidneys have visible and quantifiable stochastic vascular patterning defects with reductions in most arterial metrics during embryogenesis and postnatally. These differences in vascular metrics largely resolve in adulthood, although patterning defects are still observed. Furthermore, deletion of *Ntn1* resulted in a delay in the cessation of nephrogenesis and kidney hypoplasia. Altogether, our studies and those in the accompanying manuscript by **Luo, Gu et. al.,** which uncover similar findings, highlight the significance of netrin-1 to proper kidney vascularization and associated development.

While we find differences in vascularization in the absence of netrin-1, our analyses relate specifically to the larger arteries and arterioles of the vascular tree. SMA will only label vasculature with sufficient smooth muscle coverage, typically larger arteries and arterioles, missing the capillaries which are primarily covered by NG2+ pericytes. However, **Luo, Gu et. al.** demonstrate a lack of proper NG2+ pericyte localization in early development, suggesting netrin-1 is an important signaling molecule for all vascular mural cells. Additionally, we did not investigate venous formation and patterning during development, primarily owing to a lack of appropriate and agreed-upon markers of the venous vasculature(Daniel et al., 2018). Veins are also covered with significantly fewer SMCs due to a decreased need to regulate pressure and tone^38^. This makes SMA an unreliable venous marker at lower magnifications, such as those used by light-sheet microscopy to image the vasculature. Therefore, additional studies are necessary to determine to what extent the entire vascular plexus is disrupted in *Foxd1^GC/+^; Ntn1^fl/fl^* kidneys.

Vascular patterning and maturity are both disrupted in the *Foxd1^GC/+^; Ntn1^fl/fl^* kidneys in our study. **Luo, Gu et. al.** further define the maturity phenotype as being associated with the loss of Klf4 and accumulation of mural cells on the surface of the kidneys. We did not observe any obvious accumulation of mural cells at E15.5 or later stages, and whether Klf4 is associated with the phenotype in our studies is unknown. We did not find a difference in *Klf4* expression in our E15.5 RNA-seq, although this is 2 days later than the RNA-seq performed by **Luo, Gu et. al.** and Klf4 levels may have normalized by this time. Therefore, the observed maturation phenotypes may initiate at the earliest stages of arterialization, around E13.5, and prior to our analysis at E15.5(Daniel et al., 2018). The timing of the phenotype could explain the differences between our study and that of **Luo, Gu et. al.,** highlighting the complementary nature of our analyses. VSMCs *in vitro* express netrin receptors such as Neogenin (*Neo1*) and Unc5b and signaling through these receptors supports endothelial vSMC coverage(Lejmi et al., 2014, 2008). As such, deletion of *Ntn1* from the stroma may ultimately impact SMC maturation or recruitment in the kidney and, in turn, affect vascular patterning. This is similar to the findings of Hurtado et al. who showed that Pbx1-mediates mural cell maturation ultimately influencing proper vascular patterning within the developing kidney(Hurtado et al., 2015). Alternatively, patterning of the endothelium may influence vascular maturation. Deletion of *Unc5b* from the kidney endothelium and the assessment of vascular patterning and maturity would potentially help sort out if the patterning defects we observed are independent of vSMC coverage. The lack of significant phenotypes in the *Unc5c* knockouts suggest it is either not a major receptor for netrin-1 in the kidney, or it is functionally redundant. Considering netrin-1 can signal through a large number of receptors, including integrins, and display receptor independent cross-repressive activities with LRIG proteins, uncovering the receptor(s) and downstream signaling mechanisms of netrin-1 in the kidney may prove challenging (Abdullah et al., 2021; Abraira et al., 2008; Stanco et al., 2009; Sun et al., 2011; Yebra et al., 2003).

The kidney hypoplasia observed in our *Foxd1^GC/+^; Ntn1^fl/fl^* mutants may be linked to the relative vascular immaturity we observed. This could perturb metabolic processes within the kidney leading to downregulation of the glycolysis pathways and associated genes in our RNA-seq data, and ultimately affect proper kidney growth and prolong the nephrogenic period(Liu et al., 2017). Alternatively, a mechanism independent of the vasculature may perturb proper kidney development. Altered interactions between netrin-1 and Bmp4 could lead to the hypoplastic phenotype. Bmp4 is a negative regulator of ureteric branching through its modulation of Gdnf signaling(Miyazaki et al., 2000). Dysregulation of Gdnf signaling can result in hypoplasia and agenesis at the extreme(Moore et al., 1996; Pichel et al., 1996; Sánchez et al., 1996). Netrin-1 suppresses Bmp4 signaling *in vitro*, and netrin-1 mediated modulation of Bmp signaling within the kidney could help balance Gdnf signaling(Abdullah et al., 2021). It is possible that loss of *Ntn1* results in increases in Bmp4 and a subsequent reduction in Gdnf activity, leading to renal hypoplasia. Our RNA-seq analyses did not reveal any significant perturbations to *Bmp4* o r *Gdnf* levels, although assessing an earlier timepoint when the hypoplasia is initiated may be more informative and help sort out the cause of the hypoplastic phenotype.

Whether the development of a kidney with abnormal vascular patterns affects physiological function remains unknown and will be the focus of future studies. *Foxd1^GC/+^; Ntn1^fl/fl^* animals do live into adulthood, with no significant kidney or health phenotypes. However, if the smaller vascular beds in important functional regions of the nephron such as the glomerulus and tubules are organized properly, then physiological function may be largely unperturbed. However, injury, disease, or aging may exacerbate or induce changes in physiological function in these animals regardless. Understanding whether precise vascular patterning is critical to function, both in health and under physiological stress, will help inform efforts to vascularize tissues for functional transplant.

As the kidney lacks regenerative mechanisms, *de novo* kidney engineering and *in situ* repair using *in vitro* generated nephrons or iPSC-derived renal organoids are considered attractive therapeutic possibilities(Morizane et al., 2015; Taguchi et al., 2014; Takasato et al., 2015; Wu et al., 2018). Unfortunately, neither treatment is currently approaching clinical relevance(Bantounas et al., 2018). With current protocols, renal organoids form basic nephron structures but eventually become necrotic from a deficit of oxygen and nutrients since functional endothelium is not present(Grebenyuk and Ranga, 2019). Interestingly, cells expressing the pan-endothelial marker CD31 occur *de novo* in organoids, but without perfusion they do not form connected, functional networks, resulting in a lack of proper lumenization, flow, and patterning and their eventual regression(Homan et al., 2019; Ryan et al., 2021; Taguchi et al., 2014; Wu et al., 2018). Additionally, the vessels would need to obtain appropriate mural cell coverage to support function. Therefore, studies such as ours and those of **Luo, Gu et. al.** provide critical insights into how the renal vasculature is patterned, matures, and contributes to kidney development and will help guide kidney engineering strategies.

## Material and methods

### Animals

All animal studies were approved by the Office of Animal Care and Use at the University of North Carolina at Chapel Hill and the UNC Institutional Animal Care and Use Committee (IACUC). Procedures were performed under IACUC-approved protocols for mice (16-276.0, 19- 183.0, 22-136). Animal husbandry is performed by UNC’s Division of Comparative Medicine (DCM). All animals in this study were kept in environmental conditions consisting of temperatures between 68-74°F, with humidity ranges of 30-70%. No more than five adult animals were housed per cage, and all cages were provided a 12-hour light/dark cycle. Animals are fed LabDiet PicoLab Select Rodent 50 IF/6F (Catalog #, 5V5R). Timed matings were set up and plugs were ascertained by visual and probed inspection. The presence of a plug was considered 0.5 days gestation. The day of birth (E19.5) was considered postnatal day 0 (P0), pups born before or after E19.5 were not utilized in these studies.

All mouse lines are maintained on the C57Bl6/J background (Jackson Labs, Strain #000664). Details of genetically edited mouse lines utilized are as follows: *Foxd1^GC/+^* (Strain #012463)(Humphreys et al., 2010; Kobayashi et al., 2014) and *Ntn1^fl/fl^* (Strain #028038)(Bin et al., 2015) mice were purchased from Jackson Labs, *TrkA* knockout mice(Liebl et al., 2000) were a gift from Lino Tessarollo at the NIH National Cancer Institute Center for Cancer Research, and *Unc5c* knockout mice(Ackerman et al., 1997) were a gift from Susan Ackerman at the University of California, San Diego. *Foxd1^GC/+^; Ntn1 ^fl/fl^* mice were generated by crossing Cre-expressing *Foxd1^GC/+^; Ntn1 ^f/+^* to *Ntn1^fl/fl^* mice unless otherwise noted. Controls were either *Ntn1^c/c^* or *Foxd1^GC/+^; Ntn1 ^fl/+^* as we did not find any qualitative or quantitative differences between genotypes for any of the analyses.

*Ntn3* knockout mice were generated by the UNC Animal Models Core using CRISPR/Cas9-mediated genome editing. Guide RNAs *Ntn3*-gRNA1 (CGAGGAATCGCAGACGCGAT, Exon 1, chr17:24208777-24208796) and *Ntn3*-gRNA2 (GAAAGCGTGTCCACGAGCCG, Intron 5, chr17:24206891-24206910) were designed to delete the majority of the *Ntn3* coding region spanning Exons 1-6. Guide RNAs were generated within our lab and provided to the animal models core at estimated concentrations of 500ng/µl and 450ng/µl, respectively. Guide RNAs were prepared as in O’Brien et al.(O’Brien et al., 2018). Recombinant Cas9 protein was expressed and purified by the UNC Protein Expression Core using a plasmid provided by the Animal Models Core. Two hundred and ten fertilized embryos (zygotes) were harvested from 9 super-ovulated *Ntn1^fl/fl^* females crossed to *Foxd1^GC/+^; Ntn1 ^fl/fl^* males. Zygotes were electroporated with two different reagent concentrations diluted in OptiMEM medium (Gibco) and then implanted in pseudo-pregnant recipient females. Mix 1: 1.2 µM Cas9 protein, 47 ng/µl each guide RNA. 102 embryos electroporated; 85 embryos implanted in 4 B6D2F1 recipients. Mix2: 400 nM Cas9 protein, 25 ng/µl each guide RNA. 108 embryos electroporated; 95 embryos implanted in 4 B6D2F1 recipients. Of the resulting pups, one male containing the correct deletion as confirmed by Sanger sequencing over the deletion site was used as the founder. Brain and kidney tissue was harvested, and qPCR performed to confirm the loss of *Ntn3* transcripts in knockout animals.

Genotyping primers utilized in this study: *Ntn1* F: GGCAGTGAATGTGTTTCCGTT, *Ntn1* R: ATCGCGGGATAGTGGGTTTC; *Foxd1* WT F: CTCCTCCGTGTCCTCGTC, *Foxd1* Mut F: GGGAGGATTGGGAAGACAAT, *Foxd1* Common R: TCTGGTCCAAGAATCCGAAG; *Ntn3* Internal (WT) F: GGCCTGTGGTCTGGTTACAG, *Ntn3* Internal (WT) R: TAGCCTGGGGACTTCTGACC, *Ntn3* External (deletion) F: CACCTCCCAGCAGCAAGTAAC, *Ntn3* External (deletion) R: ACGCAGAGTAGCGGACTAGG; *TrkA* 1: TGTACGGCCATAGATAAGCAT, *TrkA* 2: TGCATAACTGTGTATTTCAC, *TrkA* 3: CGCCTTCTTGACGAGTTCTTCTG; *Unc5c* WT F: CACTCTATGGAAATGGCTGAAT, *Unc5c* WT R: GTCCTCCAATCCAAGAACTG, *Unc5c* Mut F: CAGGAGAAGATACATTTAACCAC, *Unc5c* Mut R: GACAGAAGAGCATAGCATTCAC

### Kidney size measurements

Kidney length measurements were performed using FIJI software to analyze images taken at 2x magnification on a Leica dissecting scope. Standardization and calibration of FIJI measurements were performed using a 1µm-1mm micrometer taken at the same magnification.

### In situ hybridization

*In situ* hybridizations were performed on wildtype cryosectioned E15.5 and P2 Swiss Webster (SW, Taconic) kidneys using the “Section In Situ Hybridization (SISH) on kidney sections” protocol (Krautzberger and Guo; McMahon Lab; https://www.gudmap.org/chaise/record/#2/Protocol:Protocol/RID=N-H9AC) on GUDMAP (www.gudmap.org). Digoxygenin (DIG) labeled probes were generated following the “Digoxigenin-labelled Riboprobe Synthesis from a PCR-generated DNA templates” protocol found on GUDMAP (Little Group, GUDMAP Consortium). Primers utilized for probes are as follows: *Ntn1* F:CTTCCTCACCGACCTCAATAAC, *Ntn1* (SP6) R: GCGATTTAGGTGACACTATAGTTGTGCCTACAGTCACACACC; *Ntn3* F:GGCACGCTCTCCGCTGCA, *Ntn3* (T7) R:TAATACGACTCACTATAGGGAGTGGCTGAGTAGGAGTA; *Unc5c* F:ATGAGGAAAGGTCTGAGGGCGACAG, *Unc5c* (T7) R:TAATACGACTCACTATAGGGAGAGAGTTGAAGGTACCAA; *Unc5b* F:TCAAGTGTAATGGCGAGTG, *Unc5b* (T7) R:TAATACGACTCACTATAGGGGCCTCCATTCACATAGACGA

### Section immunostaining

For all samples, kidneys were harvested at the necessary time point. Tissue was then fixed in 4% paraformaldehyde (PFA) in 1 x PBS for 15-30 minutes (embryonic/postnatal) to 24 hours (adult). Kidneys were washed 2x in PBS and stored for up to two weeks before processing. Samples were incubated in 30% sucrose in PBS overnight at 4 °C. Tissue was embedded in OCT media, frozen with 100% EtOH and dry ice, and stored at −80 °C for up to two years. Frozen tissue was sectioned at 10-12µm. Sections were air-dried at room temperature (RT) for a minimum of 10 minutes. Samples were then incubated in 1x PBS for 5 minutes to remove OCT media. Next, sections were placed in blocking buffer (PBS with 3% BSA, 1% donkey serum, and 0.25% Triton x-100) for 20 minutes at RT. The blocking buffer was removed, then sections were incubated with primary antibodies in the blocking buffer for 1 hour at RT. Sections were then rinsed 3x in wash buffer (PBS with 0.1% Triton x-100). Next, sections were incubated with appropriate Alexa Fluor conjugated secondary antibodies at 1:1000 (Invitrogen) in blocking buffer for 20 minutes. Sections were then rinsed 3x in wash buffer and mounted with Prolong Gold with DAPI (Invitrogen). Antibodies utilized and their dilution are listed in Table S4.

### Quantification of proliferation

Mice were injected intraperitoneally with 1.25 mg of EdU (10mg/ml) 1.5 hours before euthanasia and tissue collection. For analysis of proliferation, the following procedure was used. The tissue was fixed, then sectioned and stained as described previously in this report. The Click-It EdU kit was utilized to detect EdU positive cells. Images were then collected on a Zeiss 880 confocal using a 40x oil objective. The whole kidney images were created through automated tiling of the tissue section using Zen Black software. Reconstruction of the images into a single image was performed using Imaris stitcher (Bitplane). For proliferation counts, a minimum of 10 nephrogenic caps were counted per section for each genotype and age. Quantitation of proliferation for nephron progenitors was determined as the number of cells colocalized for both EdU and Six2 divided by the total number of Six2+ cells. Ureteric bud (UB) tip niches were defined as the Ecad+ cells within the UB but above the ventral end of the nephrogenic cap. Quantitation for UB proliferation was determined as the number of cells within the UB niche colocalized with Ecad and EdU divided by the total number of Ecad+ cells within the UB niche.

### Wholemount immunostaining

The wholemount immunostaining protocol was adapted from Jafree et al.(Jafree et al., 2019) and Renier et al.(Renier et al., 2014). All following steps are performed with rocking unless otherwise noted. Samples are collected and fixed in 4% PFA for 15-30 minutes (embryonic/postnatal). Tissue is washed with rocking 2x for 20 minutes at RT. Samples are then dehydrated in an increasing MeOH/H_2_O series of 25%, 50%, 75%, and 100% for 30 minutes or 1 hour, depending on tissue size. Next, tissue is bleached overnight in 5% H _2_O_2_ (from 30% stock) in 100% MeOH. Samples are then rehydrated following a reverse MeOH/H_2_O series (100%, 75%, 50%, 25%, PBS) for 30 minutes or 1 hour, depending on tissue size. Tissue is then permeabilized overnight at 4 °C in PBS with 0.2% Triton x-100, 2.3% glycine, and 20% DMSO. The permeabilization solution is removed, and blocking buffer (PBS with 0.2% triton x-100, 6% donkey serum, and 10% DMSO) is added; the tissue is then incubated for one day at RT. The blocking buffer is then removed, and primary antibody in staining solution (PBS with 0.2% Tween-20, 0.1% heparin [from 10mg/ml stock], 3% donkey serum, and 5% DMSO) is added to samples, which are then incubated for three days at 4°C, followed by washing 4-6x for 1 hour at RT in wash buffer (PBS with 0.2% Tween-20 and 0.1% heparin [from 10mg/ml stock]). The appropriate Alexa fluor labeled secondary antibody in staining solution is then added (1:250) and incubated overnight at 4 °C. Samples are washed 4-6x for 1 hour at RT in wash buffer. Samples are then immediately moved to clearing or stored at 4 °C in 1x PBS, protected from light until ready to proceed to clearing. Antibodies utilized and their dilution are listed in Table S4.

### Evans blue

The full protocol for Evans blue dye is from Honeycutt and O’Brien(Honeycutt and O’Brien, 2021) and is described here briefly. Animals are anesthetized to a surgical plane under isoflurane. Next, the chest cavity and peritoneum are cut to expose the diaphragm. 2% Evans blue dye in 0.9% NaCl/H_2_O at 3 µl/g of body weight is injected into the left ventricle with either a 25g needle or 31g insulin syringe. The dye is allowed to circulate until the exposed areas (paws, tail, chin) turn blue, roughly 2-5 minutes. The animal is subsequently euthanized, and kidney is collected, rinsed briefly in PBS several times to remove excess dye, then fixed in 4% PFA for 24 hours at 4°C. Samples are washed in PBS and stored at 4°C until clearing.

### Tissue clearing and imaging

This protocol was adapted from the iDISCO method of tissue clearing(Renier et al., 2014). First, embryonic and small samples are placed into agarose blocks (1% agarose in tris-acetate-EDTA [TAE]). Adult kidneys are not embedded. All following steps are performed with rocking unless otherwise noted. These blocks or adult kidneys are dehydrated in an increasing MeOH/H_2_O series of 25%, 50%, 75%, and 100% for 30 minutes or 1 hour at RT, depending on tissue size. Samples are then incubated in 66% dichloromethane (DCM) / 33% MeOH for 3 hours at RT. Next, two rounds of incubation with 100% DCM at RT are performed. DCM is removed, and 100% dibenzyl ether (DBE) is added. Samples are incubated, with no rocking, until clear (1 – 24 hours) and stored indefinitely in 100% DBE at either RT or 4 °C, protected from light, until ready to proceed to imaging. Light-sheet imaging of whole kidneys was performed using a LaVision UltraMicroscope II (LaVision BioTec). 3D reconstruction of images and vascular network quantification was conducted using Imaris imaging software (Bitplane).

### Quantification of wholemount 3D vasculature

To conduct quantification of renal vasculature within Imaris, a surface of the 3D image was first created to digitally render the vasculature. Next, a binary mask of the rendered surface was created using the “Mask channel" function with “Constant inside/outside” settings for voxel intensity set to 0 outside and 100 inside. This was necessary to render the vasculature effectively solid, so that the filament tracer module within Imaris could be implemented to develop wireframes for statistical analysis. Kidneys were batched with similar parameters per age group. Then supervised automatic tracing was performed to ensure that automatically generated paths were not duplicated, or artifacts inaccurately rendered as vasculature. As both kidneys from the same animal have highly similar statistical values, with little deviation when averaged, for most studies, only one kidney per animal was needed for quantification. This method was used to quantify renal arterial trees for Evans blue in adults and SMA and CD31 staining in the embryonic and neonatal kidneys. Cortical arterioles and vascular beds were excluded from analysis.

For graphical display of vascular patterning and imaging that were not quantified, a surface was created for the desired channel. This surface was then masked using the same procedure, but “Constant inside/outside” settings for voxel intensity were set to 0 outside with the inside setting left unchecked.

### RNA-seq

Whole kidneys were collected at E15.5 from 3 separate *Ntn1^fl/fl^ x Foxd1 ^GC/+^;Ntn1^fl/+^* crosses. Three kidneys were collected of each genotype: *Foxd1^GC/+^;Ntn1^fl/+^* (heterozygous), *Foxd1^GC/+^;Ntn1^fl/fl^* (homozygous), and *Ntn1^fl/fl^* (wildtype). Kidneys were placed in RNAlater (Thermo Fisher) until all samples were collected. RNA was extracted with TRIzol (Invitrogen). RNA was sent to Cancer Genetics Inc (Morrisville, NC) for sequencing and data processing. RNA samples were converted to sequencing libraries using the TruSeq Stranded mRNA kit (Illumina Cat. #20020595). Resulting libraries were assessed for quality via analysis on a 2100 Bioanalyzer (Agilent Technologies) and quantitated by qPCR (KAPA Library Quantification; Roche: KK4824, Cat# 07960140001). Normalized and pooled libraries were sequenced on the Illumina Nextseq 550 system. Alignment and analysis were performed following a modified Tuxedo workflow. Briefly, alignment was done with STAR aligner followed by cufflinks/cuffmerge for assembled transcripts. FPKM was calculated with CuffDif and analyzed with the R package. Heterozygous samples were used as a control and comparison to the homozygous samples to account for any effects of the hemizygous *Foxd1* allele due to the knock-in of GFP-Cre into the *Foxd1* locus. A count of 5 FPKM was considered minimum criteria for expression in the whole kidney RNA-seq, genes with less than this FPKM (unless downregulated compared to the reference) were excluded from gene expression analyses. Up- and downregulated gene lists were pasted into the Database for Annotation, Visualization and Integrated Discovery (DAVID) to obtain Gene Ontology information and enrichment.

## Acknowledgments

We thank members of the O’Brien lab for technical help and critical discussions, Dr. Pablo Ariel, director of the UNC Microscopy Services Laboratory, for assistance with light-sheet imaging and Imaris rendering/analysis, Dr. Robert Currin (retired) of the UNC Hooker Imaging Core for confocal microscopy imaging assistance, and Dr. Dale Cowley, director of the UNC Animal Models Core, for help with *Ntn3* knockout mouse generation. We thank Drs. Lino Tessarollo (NCI) and Susan Ackerman (UCSD) for their kind gifts of *TrkA* and *Unc5c* knockout mice, respectively.

## Author Contributions

LLO conceived and designed the project, performed experiments, analyzed data, and helped write the manuscript. SEH helped design the experiments, performed experiments, analyzed data, and wrote the manuscript. P-EYN, DMH, YX, and SLC performed experiments and analyzed data.

