## Supplemental Figures and Legends for "Netrin-1 directs vascular patterning and maturity in the developing kidney"

### Supplemental Figure 1

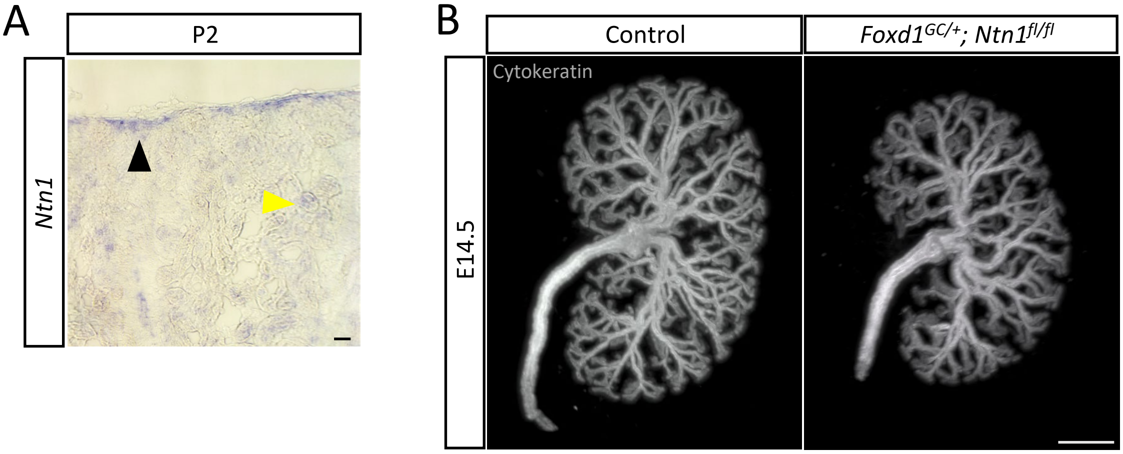

### Supplemental Figure 2

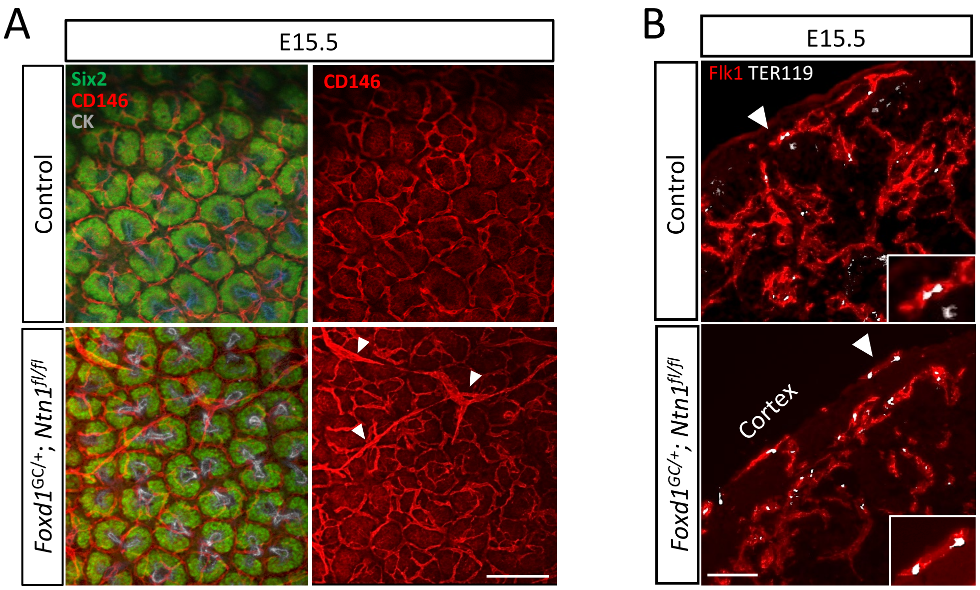

### Supplemental Figure 3

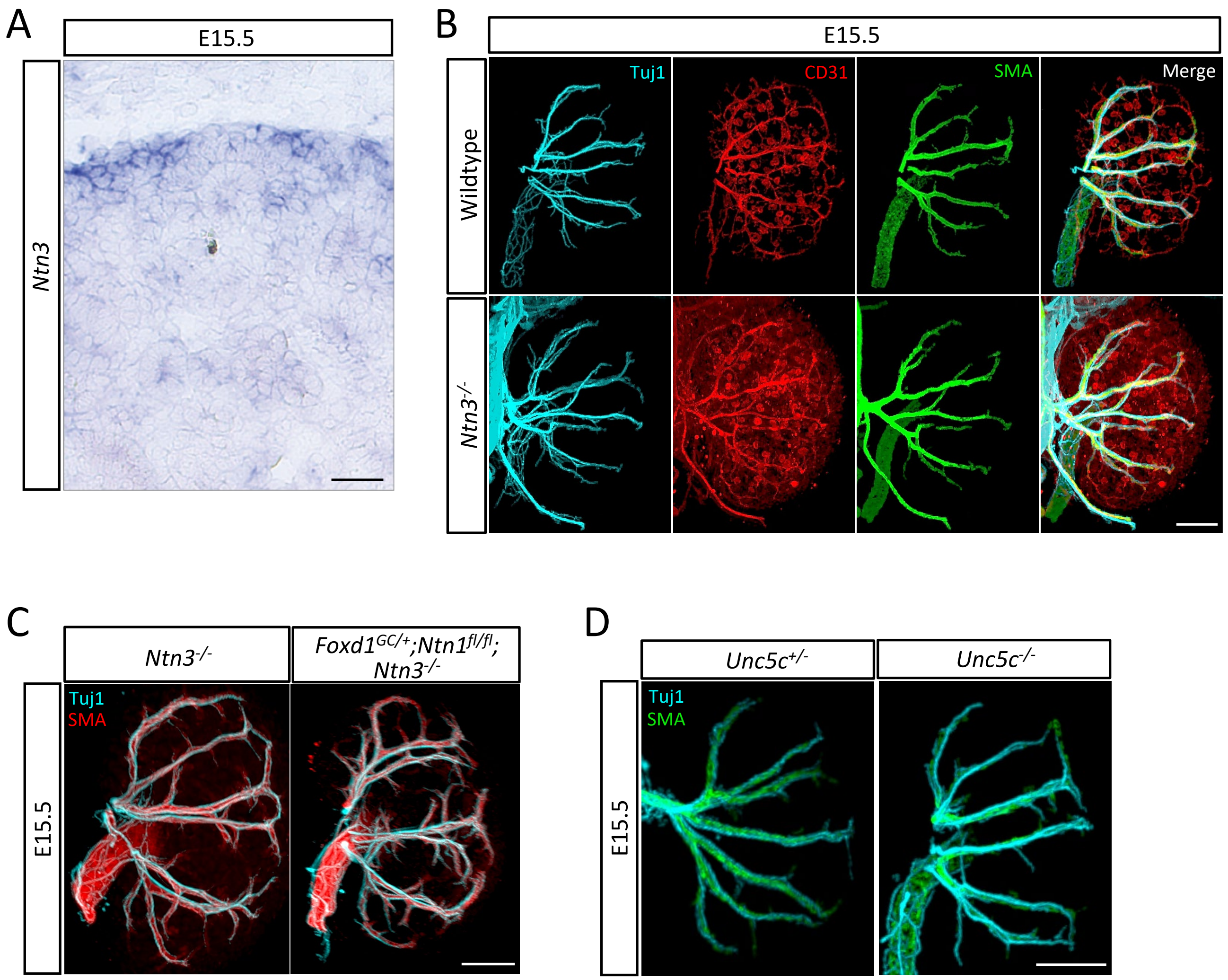

### Supplemental Figure 4

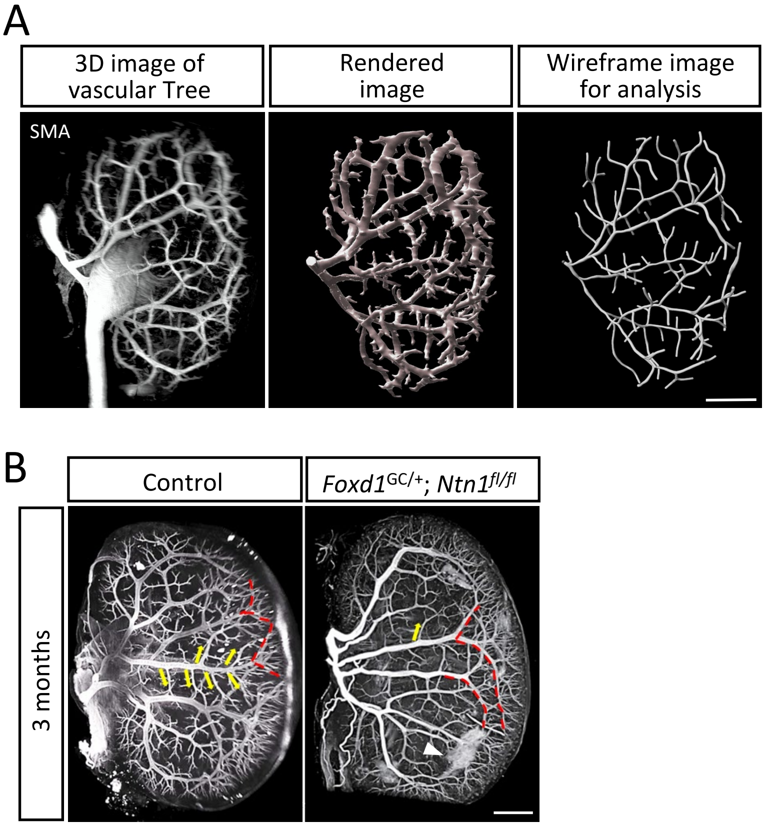

#### Supplemental Figures

**Supplemental Figure 1: *Ntn1* is expressed by postnatal stromal progenitors and deletion of *Ntn1* from Foxd1+ progenitors results in hypoplastic kidneys**

A) *In situ* hybridization of sectioned postnatal day (P)2 kidneys shows *Ntn1* mRNA expression by stromal progenitors (black arrowhead) and to a lesser extent by epithelial structures (yellow arrowhead). Scale bar=50 μm. B) Wholemount immunostaining of the ureteric tree (cytokeratin, grey) at E14.5 shows that *Foxd1*$^{GC/+}$*; Ntn1*$^{fl/f}$ kidneys are smaller than controls at this stage. Scale bar=200 μm.

**Supplemental Figure 2: *Foxd1*$^{GC/+}$*; Ntn1*$^{fl/fl}$ kidneys display vascular patterning defects around nephron progenitor niches and these vessels contain erythroid cells**

A) Confocal images of the wholemount immunostained kidney surface at E15.5 showing ectopic vessels (CD146+, red) next to nephron progenitor niches (Six2+, green) in *Foxd1*$^{GC/+}$*; Ntn1*$^{fl/fl}$ kidneys. Scale bar=100 μm. B) Sections of E15.5 kidneys show Ter119+ erythroid cells (white) are present in the vasculature (Flk1, red) of both control and *Foxd1*$^{GC/+}$*; Ntn1*$^{fl/fl}$ kidneys. Insets show higher magnification views. Scale bar=50 μm.

**Supplemental Figure 3: Deletion of *Ntn3* or *Unc5c* does not cause significant vascular patterning defects nor exacerbate the *Foxd1*$^{GC/+}$*; Ntn1*$^{fl/fl}$ phenotype with co-deletion of *Ntn3***

A) *In situ* of *Ntn3* at E15.5 on wildtype sections. A similar pattern to *Ntn1* is observed. Scale bar=50 μm. B) Wholemount immunofluorescent images of *Ntn3* knockout kidneys showing no significant differences in vascular (SMA, green; CD31, red) or neuronal (Tuj1, cyan) patterning. Scale bar=300 μm. C) Wholemount immunofluorescent images show that double knockout *Foxd1*$^{GC/+}$*; Ntn1*$^{fl/fl}$*;Ntn3*$^{-/-}$ kidneys do not show any exacerbated vascular patterning phenotypes (SMA, red;

Tuj1, cyan). Scale bar=400 μm. D) Wholemount immunofluorescent images show that vascular patterning is largely normal in *Unc5c* knockouts (Tuj1, cyan; SMA, green). Scale bar=300 μm.

**Supplemental Figure 4. Depictions of rendering used for vascular (arterial tree) analysis and how the patterning defects persist at 3 months**

A) Micrographs showing the original image of vascular trees, the Imaris rendered image, and resulting wireframe image used for analysis. Scale bar=500 μm B) Images of 3-month-old adult kidneys stained with Evans blue showing the persistence of vascular patterning defects such as meandering vessels (red dotted lines) and reduced branching (yellow arrows). Clouds of Evans blue are occasionally observed (white arrowhead), potentially representing vascular leakage. Scale bar=1000 μm.

#### Supplemental Tables

**Table S1:** RNA-seq data for WT (*Ntn1*$^{fl/fl}$), HET (*Foxd1*$^{GC/+}$;*Ntn1*$^{fl/+}$) and MUT (*Foxd1*$^{GC/+}$;*Ntn1*$^{fl/fl}$ ) kidneys

**Table S2:** Genes upregulated $\geq$ 1.2-fold and the associated Gene Ontology (GO) Biological Processes

**Table S3:** Genes downregulated $\geq$ 1.2-fold and the associated Gene Ontology (GO) Biological Processes

**Table S4:** List of antibodies utilized for section and wholemount immunostaining

#### Supplemental Videos

**Supplemental Video 1:** Video associated with Fig. 3C showing vascular patterning of a control kidney. Tuj1+ nerves: cyan; CD31+ endothelium: red

**Supplemental Video 2:** Video associated with Fig. 3C showing vascular patterning of a *Foxd1*$^{GC/+}$;*Ntn1*$^{fl/fl}$ kidney. Tuj1+ nerves: cyan; CD31+ endothelium: red

**Supplemental Video 3:** Video associated with Fig. 4B showing vascular patterning of a control kidney. Tuj1+ nerves: cyan; CD31+ endothelium: red, SMA+ mural cells: green

**Supplemental Video 4:** Video associated with Fig. 4B showing vascular patterning of a *Foxd1*$^{GC/+}$;*Ntn1*$^{fl/fl}$ kidney. Tuj1+ nerves: cyan; CD31+ endothelium: red, SMA+ mural cells: green
